# The draft genome sequence of herbaceous diploid bamboo *Raddia distichophylla*

**DOI:** 10.1101/2020.04.27.064089

**Authors:** Wei Li, Cong Shi, Kui Li, Qun-jie Zhang, Yan Tong, Yun Zhang, Jun Wang, Lynn Clark, Li-zhi Gao

**Author notes:** These authors contributed equally to this work (Wei Li, Cong Shi, Kui Li). Correspond author: E-mail address &.

## Abstract

Bamboos are important non-timber forest plants widely distributed in the tropical and subtropical regions of Asia, Africa, America, and Pacific islands. They comprise the Bambusoideae in the grass family (Poaceae), including approximately 1,700 described species in 127 genera. In spite of the widespread uses of bamboo for food, construction and bioenergy, the gene repertoire of bamboo still remains largely unexplored. *Raddia distichophylla* (Schrad. ex Nees) Chase, belonging to the tribe Olyreae (Bambusoideae, Poaceae), is diploid herbaceous bamboo with only slightly lignified stems. In this study, we report a draft genome assembly of the approximately ∼589 Mb whole-genome sequence of *R. distichophylla* with a contig N50 length of 86.36 Kb. Repeated sequences account for ∼49.08% of the genome, of which LTR retrotransposons occupy ∼35.99% of whole genome. A total of 30,763 protein-coding genes were annotated in the *R. distichophylla* genome with an average transcript size of 2,887 bp. Access to this herbaceous bamboo genome sequence will provide novel insights into biochemistry, molecular-assisted breeding programs and germplasm conservation for bamboo species world-wide.

## INTRODUCTION

Bamboos are important non-timber forest plants with a wide native geographic distribution in tropical, subtropical and temperate regions except for Europe and Antarctica (BambooPhylogenyGroup, 2012). Bamboos are of notable economic and cultural significance worldwide, and can be used as food, bioenergy and building materials. Bamboos comprise the Bambusoideae in grass family (Poaceae), including approximately 1,700 described species in 127 genera (Clark and Oliveira, 2018; Soreng et al., 2017; Vorontsova et al., 2016). Molecular phylogenetic analysis shows that Bambusoideae falls into the Bambusoideae-Oryzoideae-Pooideae (BOP) clade and is phylogenetically sister to Pooideae (Saarela et al., 2018). Bambusoideae may be divided into two morphologically distinct growth forms: woody bamboos and herbaceous bamboos (tribe Olyreae); woody bamboos can be further divided into two lineages: temperate woody (Arundinarieae) and tropical woody (Bambuseae) (BambooPhylogenyGroup, 2012; Kelchner and Group, 2013; Sungkaew et al., 2009).

Although the phylogenetic relationship between the two woody bamboo lineages is still controversial (BambooPhylogenyGroup, 2012; Bouchenak-Khelladi et al., 2008; Clark et al., 2015; GrassPhylogenyWorkingGroup et al., 2001; Sungkaew et al., 2009; Triplett et al., 2014; Wysocki et al., 2016), their shared phenology and sexual systems are suggestive of common ancestry (Wysocki et al., 2016). Woody bamboos form highly lignified, usually hollow culms, complex rhizome systems, specialized culm leaves, complex vegetative branching, and outer ligules on the foliage leaves (BambooPhylogenyGroup, 2012; Clark et al., 2015). Woody bamboos possess bisexual flowers and flower at intervals between 7 and 120 years usually followed by a die-off (BambooPhylogenyGroup, 2012; Dransfield and Widjaja, 1995; Judziewicz et al., 1999). All woody bamboos with known chromosome counts are uniformly polyploidy (Soderstrom, 1981).

The tribe Olyreae comprises 22 genera and 124 described species native to tropical America except *Buergersiochloa* and *Olyra latifolia* (BambooPhylogenyGroup, 2012; Clark et al., 2015). Herbaceous bamboos are characterized by usually weakly developed rhizomes and less lignification in the culms. Culm leaves and foliage leaves with the outer ligule are absent in herbaceous bamboos. In contrast to woody bamboos, herbaceous bamboos have at least functionally unisexual flowers and they flower annually or seasonally for extended periods (Clark et al., 2015; Gaut et al., 1997; Kelchner and Group, 2013; Oliveira et al., 2014; Wysocki et al., 2015). The tribe is fundamentally diploid, but chromosome counts indicating tetraploidy or hexaploidy or possibly even octoploidy (in *Eremitis* genus) are available (Judziewicz et al., 1999; Soderstrom, 1981).

Since the first comparative DNA sequence analysis of bamboos by Kelchner and Clark (Kelchner and Clark, 1997), a number of studies have been further carried out combined various biotechnologies (Das et al., 2005; Gui et al., 2010; Oliveira et al., 2014; Peng et al., 2013; Sharma et al., 2008; Sungkaew et al., 2009; Wysocki et al., 2016; Zhang et al., 2011). Peng et al. (Peng et al., 2013) reported the first draft genome of tetraploid moso bamboo (*Phyllostachys edulis*). Recently, Guo et al. (Guo et al., 2019) have released four draft genomes of bamboo lineages, including *O. latifolia, R. guianensis, Guadua angustifolia*, and *Bonia amplexicaulis*. The lack of a high-quality genome sequence for a diploid bamboo has been an impediment to our understanding of bamboo biology and evolution.

*R. distichophylla* (Schrad. ex Nees) Chase, a diploid herbaceous bamboo, is almost completely restricted to the forests of eastern Brazil (Oliveira et al., 2014). In this study, we generate the complete genome sequence of *R. distichophylla* through the Illumina sequencing platform. The availability of the fully sequenced and annotated genome will provide functional, ecological and evolutionary insights into the bamboo species.

## METHODS AND MATERIALS

### Sample collection, total DNA and RNA extraction and sequencing

The source plant was an individual of *R. distichophylla* grown in cultivation at the R.W. Pohl Conservatory, Iowa State University. Fresh and healthy leaves were harvested and immediately frozen in liquid nitrogen, followed by storage at −80°C in the laboratory prior to DNA extraction. A modified CTAB method (Porebski et al., 1997) was used to extract high-quality genomic DNA. The quantity and quality of the extracted DNA were examined using a NanoDrop D-1000 spectrophotometer (NanoDrop Technologies, Wilmington, DE) and electrophoresis on a 0.8% agarose gel, respectively. A total of 9 paired-end sequencing libraries, spanning 180, 300, 500, 2 000, 5 000, 10 000 and 20 000 bp, were prepared and sequenced on Illumina HiSeq 2000 platform.

Total RNA was extracted from four tissues (root, stem, young leaf and female inflorescence), using a Water Saturated Phenol method. RNA libraries were built using the Illumina RNA-Seq kit (mRNA-Seq Sample Prep Kit P/N 1004814). The extracted RNA was quantified using NanoDrop-1000 UV-VIS spectrophotometer (NanoDrop), and RNA integrity was checked using Agilent 2100 Bioanalyzer (Agilent Technologies, Palo Alto, CA, USA). For each tissue, only total RNAs with a total amount ≥ 15 μg with a concentration ≥ 400 ng/μl, RNA integrity number (RIN) ≥ 7, and rRNA ratio ≥ 1.4 were used for constructing cDNA library according to the manufacturer’s instruction (Illumina, USA). The libraries were then sequenced (100 nt, pair-end) with Illumina HiSeq 2000 platform.

### *De novo* assembly of the *R. distichophylla* genome

Two orthogonal methods were used to estimate the genome size of *R. distichophylla*, including *k*-mer frequency distribution and flow cytometric analysis. Firstly, we generated the 17-mer occurrence distribution of sequencing reads using GCE v1.0.0 (Liu et al., 2013), and the genome size was then calculated with the equation 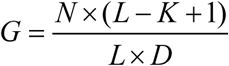, where *G* represents the genome size; *N* is the total number of reads used; *L* is the average length of reads; *K* is set to 17. Secondly, the genome size was further estimated and validated using flow cytometry analysis. We employed the rice cultivar *Nipponbare* as an inner standard with an estimated genome size of 389 Mb (IRGSP., 2005).

Paired-end sequencing reads were processed to remove adaptor and low quality sequences using Trimmomatic v0.33 (Bolger et al., 2014). Reads were retained only if both paired reads passed quality control filtering. We assembled the WGS sequencing reads using Platanus v1.2.1 software (Rei Kajitani, 2014), which is optimized for highly heterozygous diploid genomes. First, the high-quality paired-end Illumina reads from short-insert size libraries (⩽ 500 bp) were assembled into contig sequences using Platanus. The assembled contigs were then scaffolded using SSPACE v3.0 (Boetzer et al., 2011) using Illumina mate pair data. GapCloser v1.12 (Li et al., 2010) was used to fill gaps within the scaffolds.

The three methods were employed to assess the quality and completeness of the *R. distichophylla* genome. First, high-quality reads from short-insert size libraries were mapped to the genome assembly using BWA v0.7.15 (Li and Durbin, 2009). Second, the genome assembly was checked with benchmarking universal single-copy orthologs (BUSCO) (Simao et al., 2015). Third, the RNA sequencing reads generated in this study were assembled using Trinity v r20131110 (Grabherr et al., 2011), the assembled transcripts were then aligned back to the assembled genome using GMAP v2014-10-2 (Wu and Watanabe, 2005) at 30% coverage and 80% identity thresholds.

### Annotation of repetitive sequences and non-coding RNA genes

We used a combination of homology-based and *de novo* approaches to identity the repetitive sequences in the *R. distichophylla* genome. RepeatModeler v1.0.10 (Tarailo-Graovac and Chen, 2009), which included two *de novo* repeat finding programs, RECON (Bao and Eddy, 2002) and RepeatScout (Price et al., 2005), was used for the construction of the repeat library. This produced library, along with the Poaceae repeat library, were used as the reference database for RepeatMasker (Tarailo-Graovac and Chen, 2009). Simple sequence repeats (SSRs) were identified in the genome sequence using the MISA perl script (Thiel et al., 2003) with the default settings: monomer (one nucleotide, n ⩾ 12), dimer (two nucleotides, n ⩾ 6), trimer (three nucleotides, n ⩾ 4), tetramer (four nucleotides, n ⩾ 3), pentamer (five nucleotides, n ⩾ 3), and hexamer (six nucleotidess, n ⩾ 3).

Non-coding RNAs genes play important roles in many cellular processes. The five different types of non-coding RNA genes, namely transfer RNA (tRNA) genes, ribosomal RNA (rRNA) genes, small nucleolar RNA (snoRNAs) genes, small nuclear RNA (snRNAs) genes and microRNA (miRNAs) genes, were predicted using various *de novo* and homology search methods. We used tRNAscan-SE algorithms (version 1.23) (Lowe and Eddy, 1997) with default parameters to identify the tRNA genes. The rRNA genes (8S, 18S, and 28S), which is the RNA component of the ribosome, were predicted by using RNAmmer algorithms (v1.2) (Lagesen et al., 2007) with default parameters. The snoRNA genes were annotated using snoScan v1.0 (Lowe and Eddy, 1999) with the yeast rRNA methylation sites and yeast rRNA sequences provided by the snoScan distribution. The snRNA genes were identified by INFERNAL software (v1.1.2) (Nawrocki et al., 2009) against the Rfam database (release 9.1) with default parameters.

### Genome annotation

The gene prediction pipeline combined the *de novo* method, the homology-based method and the EST-aided method. Augustus v2.5.5 (Stanke et al., 2004) and Fgenesh (Salamov and Solovyev, 2000) were used to perform the *de novo* prediction. The protein sequences of moso bamboo, stiff brome, barley, maize, *Oropetium thomaeum*, foxtail millet, rice and sorghum were mapped to the genome using Exonerate (Slater and Birney, 2005). To further aid the gene annotation, Illumina RNA-seq reads were assembled using the Trinity software (v20131110) (Grabherr et al., 2011). The resulting transcripts were then aligned to the soft-masked genome assembly using GMAP v2014-10-2 (Wu and Watanabe, 2005) and BLAT v35 (Kent, 2002). The potential gene structures were derived using PASA v20130907 (Program to Assemble Spliced Alignments) (Haas et al., 2003). All gene models produced by the *de novo*, homology-based and EST-aided methods were integrated using GLEAN (Elsik et al., 2007).

The predicted genes were searched against Swiss-Prot database (Boeckmann et al., 2003) using BLASTP (e-value cutoff of 10^−5^). The motifs and domains within gene models were identified by InterProScan (Jones et al., 2014). Gene Ontology terms and KEGG pathway for each gene were retrieved from the corresponding InterPro entry. Gene functions were also assigned to TrEMBL (Boeckmann et al., 2003) database using BLASP with an e-value threshold of 10^−5^.

### Data availability

All sequencing reads have been deposited in the NCBI Sequence Read Archive SRR8759078 to SRR8759084 (2019) and BIG Genome Sequence Archive CRR049770 to CRR049776 (2019).The assembled genome sequence is available at the NCBI and BIG Genome Warehouse under accession number SPJY00000000 and GWHAAKD00000000, respectively. Table S1 shows the whole genome sequencing (WGS) reads used to assemble the *R. distichophylla* genome. Table S2 shows the summary of RNA sequencing (RNA-Seq) of *R. distichophylla*. Table S3 shows the estimation of genome size based on *K*-mer analysis. Table S4 shows the summary of genome assembly. Table S5 shows the validation of the *R. distichophylla* genome assembly using reads mapping BUSCO, and transcript alignments. Table S6 shows the statistics of repeat sequences in the *R. distichophylla* and moso bamboo genomes. Table S7 shows the summary of types and number of simple sequence repeats in the *R. distichophylla* and moso bamboo genomes. Table S8 shows the non-coding RNA genes in the *R. distichophylla* genome. Table S9 shows the statistics of predicted protein-coding genes in the *R. distichophylla* genome. Table S10 shows the functional annotation of the *R. distichophylla* protein-coding genes.

## RESULTS AND DISCUSSION

We performed whole-genome sequencing with the Illumina sequencing platform. A total of 253.94 Gb short sequencing reads were generated (∼272.89-fold coverage) (**Table S1**). A total of 21.99 Gb RNA-seq data was obtained from root, stem, young leaf and female inflorescence (**Table S2**). Based on the *K*-mer analysis, we estimated the genome size of *R. distichophylla* to be ∼608 Mb (**Table S3, Fig. 1**). Flow cytometry analysis estimated the genome size of *R. distichophylla* to be ∼589 Mb, which is close to the obtained result from the *k*-mer analysis. The final assembly amounted to ∼580.85 Mb, 95.56% of the estimated genome size. The N50 lengths of the assembled contigs and scaffolds were ∼86.36 Kb and ∼1.81 Mb, respectively (**Table 1, Table S4**).

**Table 1.**
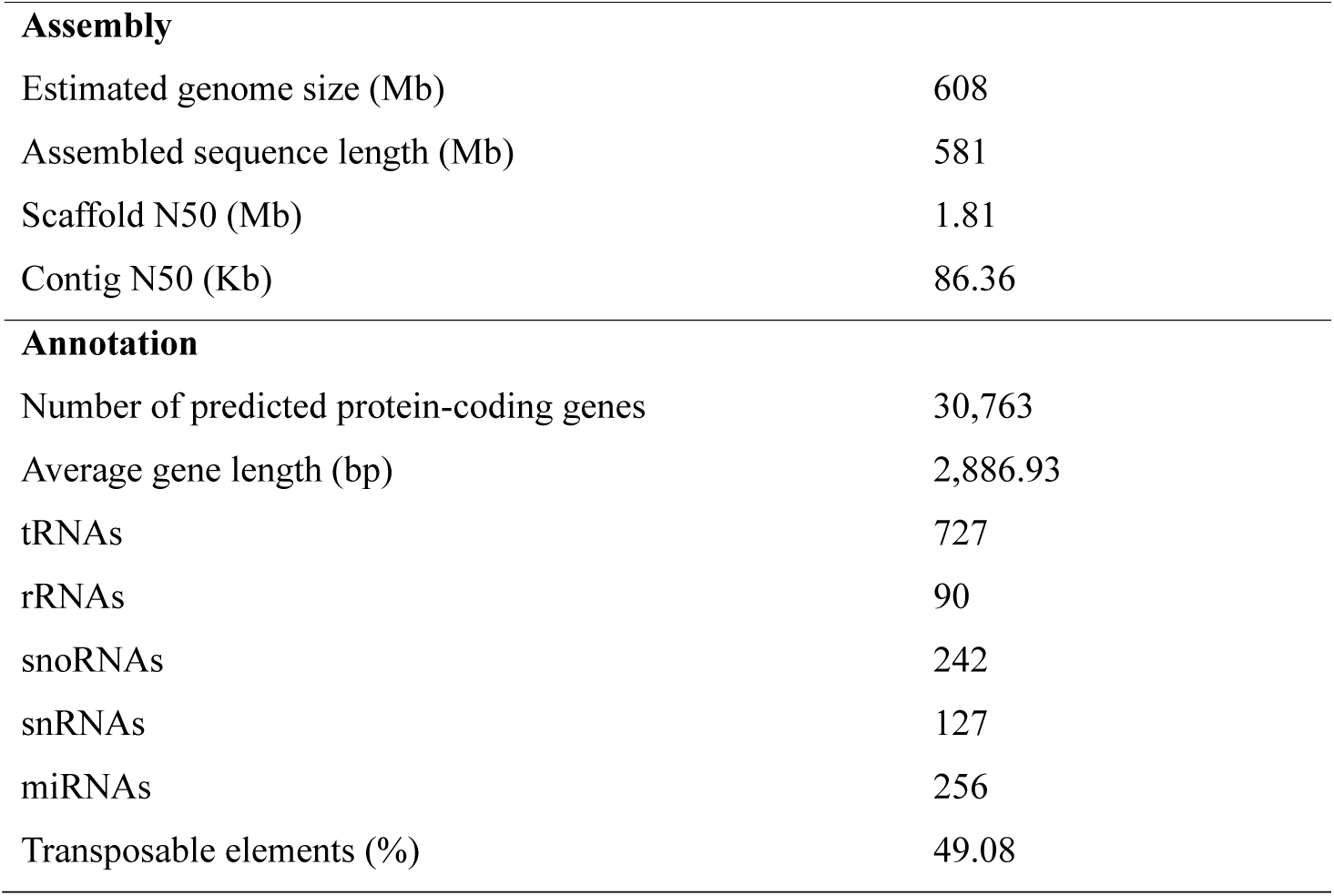
Summary of the genome assembly and annotation of *R. distichophylla*.

**Figure 1.**
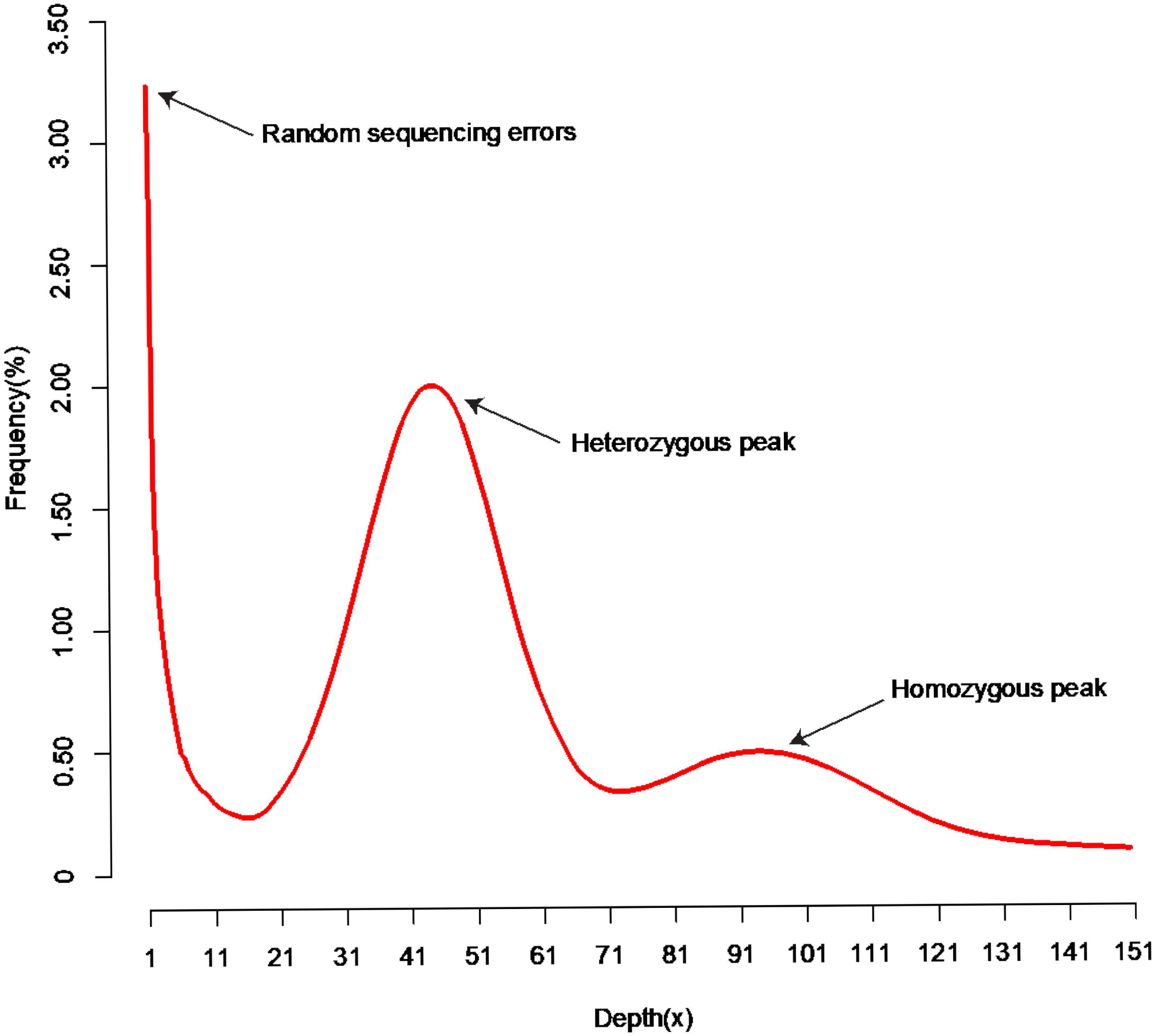
The 17-mer distribution of sequencing reads from *R. distichophylla*. The occurrence of 17-mer was calculated using GCE based on the sequencing data from short insert size libraries (insert size ≤ 500 bp) of *R. distichophylla*. The sharp peak on the left with low depths represents the essentially random sequencing errors. The middle and right peaks indicate the heterozygous and homozygous peaks, the depths of which are 52 and 103, respectively.

To validate the genome assembly quality, we first mapped ∼211 M of high-quality reads to the genome sequences. Our results revealed that nearly 89.25% Illumina reads were mapped to the genome assembly (**Table S5**); second, BUSCO was used to assess the completeness of the genome assembly. The percentage of completeness for our assembly was 92.08 in the Embryophyta lineage (**Table S5**); and finally, we mapped the assembled transcripts to the genome sequences. Approximately 78.04% of the transcripts could be mapped to the genome (**Table S5**).

The annotation of repeat sequences showed that approximately 49.08% of the *R. distichophylla* genome consists of transposable elements (TEs), lower than the amount (63.15%) annotated in the moso bamboo genome (Peng et al., 2013) with the same methods (**Table S6, Fig. 2**). LTR retrotransposons were the most abundant TE type, occupying roughly 35.99% of the *R. distichophylla* genome. In total, 220,737 and 496,819 SSRs were found in the *R. distichophylla* and moso bamboo genome, respectively, with trimer and tetramer as the most abundant SSR types (**Table S7**). Among the trimer motifs, (CCG/GGC)n were the predominant repeat in *R. distichophylla*, whereas (CCG/GGC)n, (AGG/CCT)n and (AAG/CCT)n showed a similar proportions in the moso bamboo genome (**Fig. 3**). The identified SSRs will provide valuable molecular resources for germplasm characterization and genomics-based breeding programs. In total we identified 727 tRNA genes, 90 rRNA genes, 242 snoRNA genes, 127 snRNA genes, and 256 miRNA genes, respectively (**Table 1, Table S8**).

**Figure 2.**
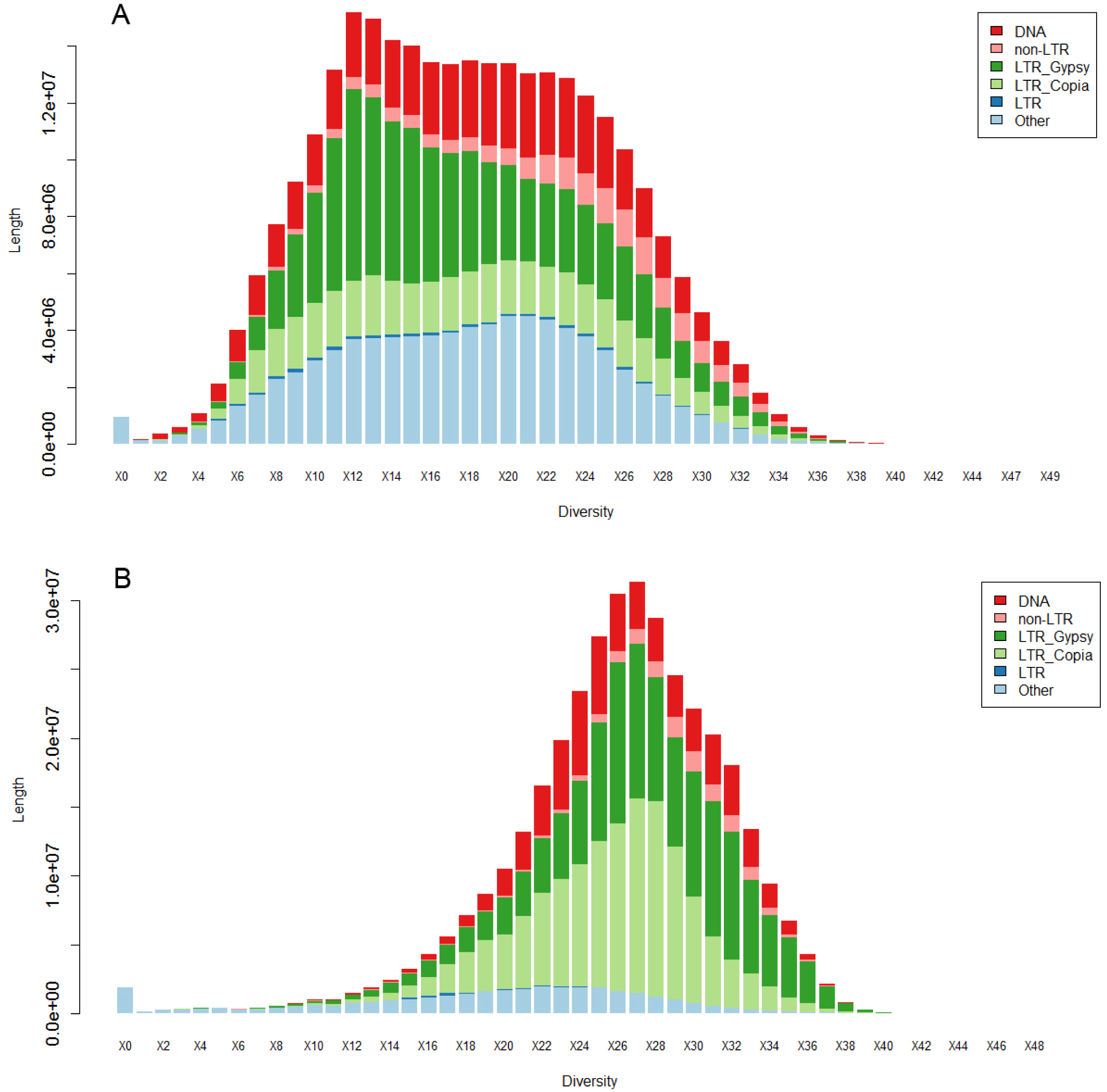
Evolutionary history of TE super-families in the *R. distichophylla* and moso bamboo genomes. A) The occupied TE lengths in *R. distichophylla* genome; B) the occupied TE lengths in the moso bamboo genome. The divergence was calculated between TEs and the consensus sequences generated by RepeatMasker.

**Figure 3.**
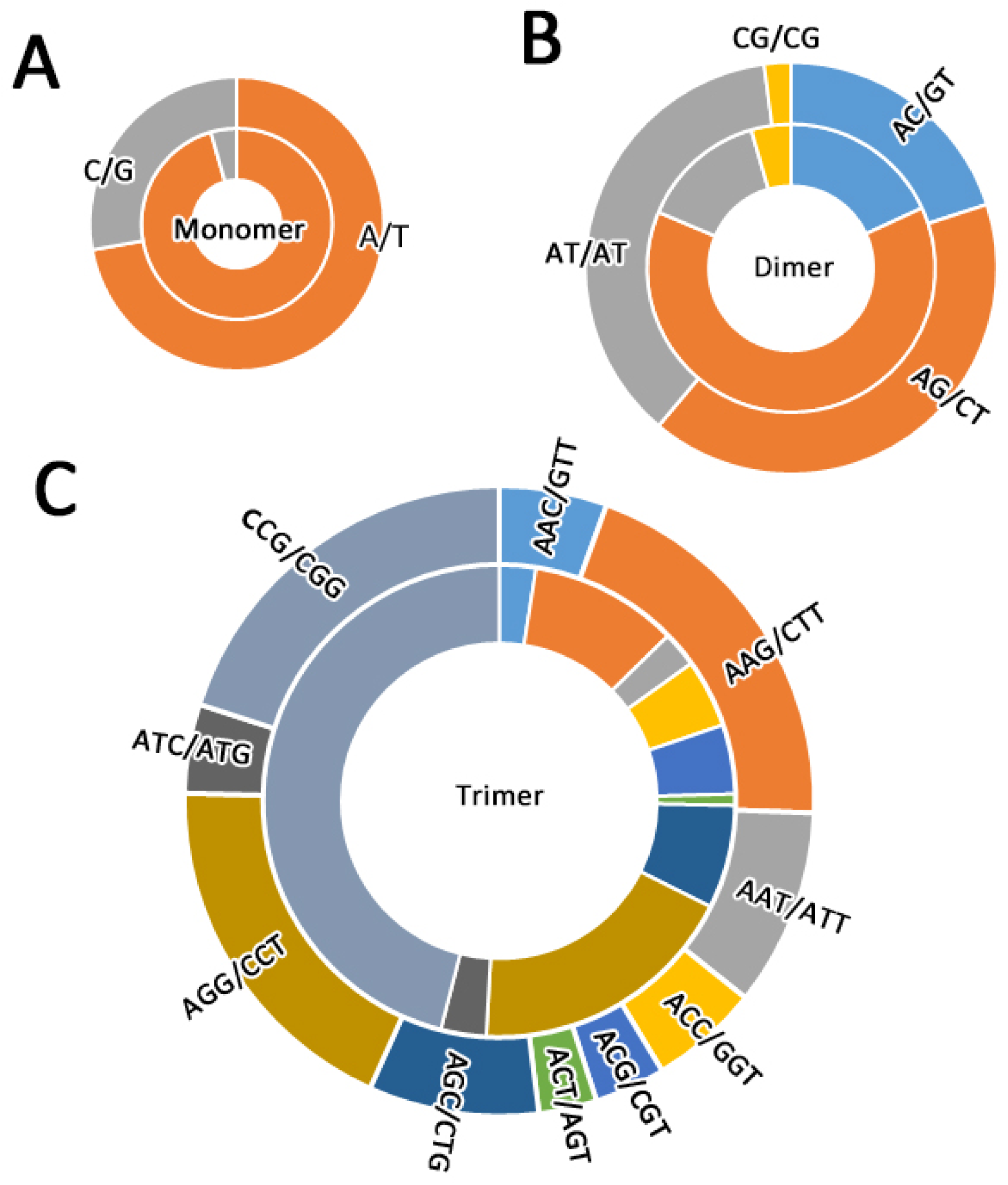
Distribution of different SSR motifs in the *R. distichophylla* and moso bamboo genomes. The outer circle represents moso bamboo and the inner circle denotes *R. distichophylla*. A) monomer distribution in the two genomes; B) dimer distribution in the two genomes; C) trimer distribution in the two genomes.

In combination with *ab initio* prediction, protein and expressed sequence tags (ESTs) alignments, we generated a gene set consisted of 30,763 protein-coding genes (**Table 1, Table S9**), with an average length of 2,887 bp and an average coding sequence length of 1,099 bp (**Table S9, Fig. 4**). Among these genes, 88.85% had significant similarities to sequences in the public databases (**Table S10**).

**Figure 4.**
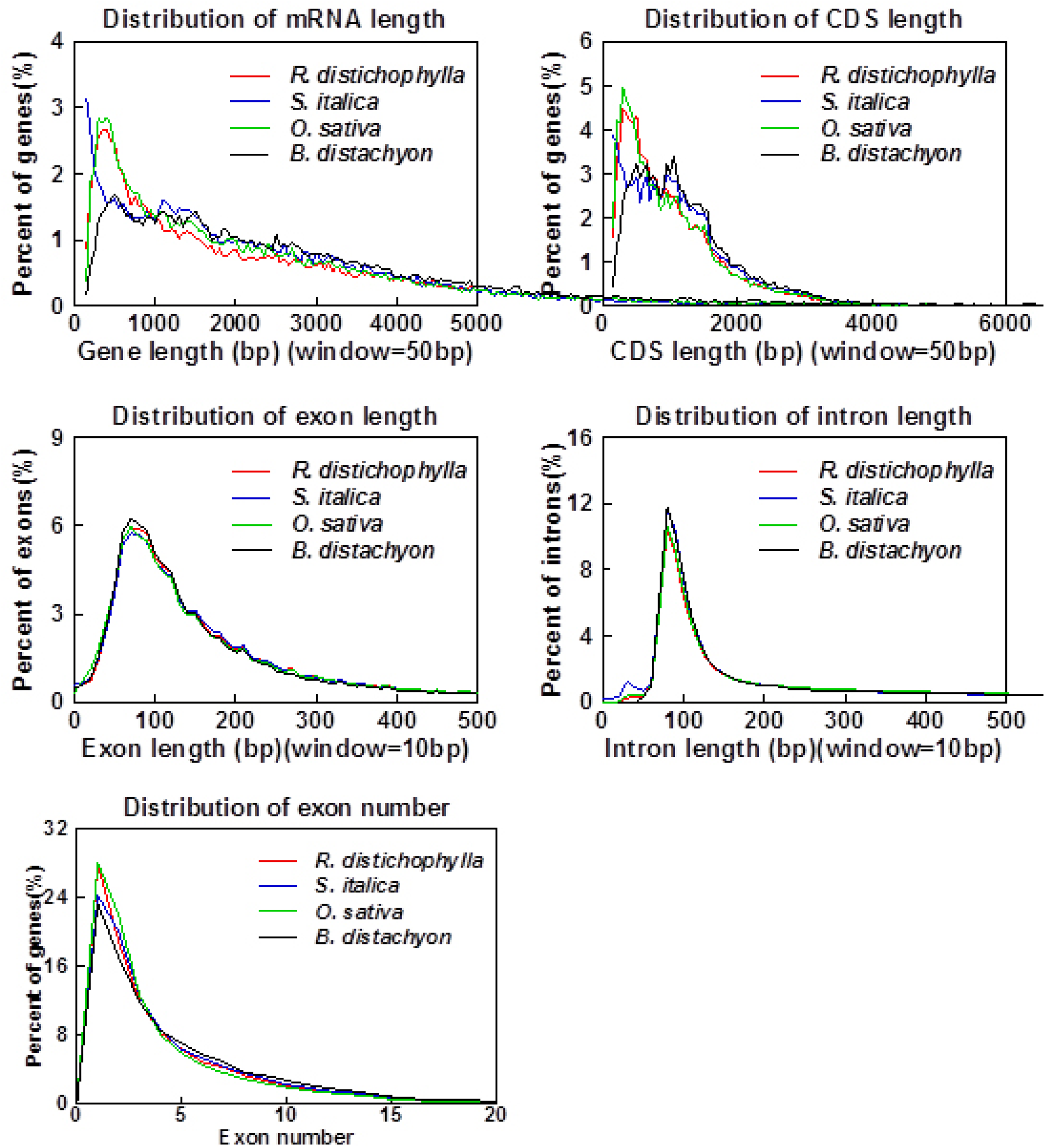
Comparisons of gene features among *R. distichophylla* and three other plant species. Gene features include mRNA length, CDS length, exon length, intron length and exon number. No obvious length differences of gene features were observed among these three plant species (*Setaria italica, Brachypodium distachyon* and *Oryza sativa*).

## ACKNOWLEDGEMENTS

This work was supported by Yunnan Innovation Team Project and the start-up grant from South China Agricultural University (to L. G.).

## Supplementary Tables

**Supplementary Table 1.** Whole genome sequencing (WGS) reads used to assemble the *R. distichophylla* genome.

| **Data Type** | **Lane Information** | **Insert Size (bp)** | **Read Length (bp)** | **Raw Data (Gb)** | **Clean Data (Gb)** | **Sequence Coverage (×)\*** |
| --- | --- | --- | --- | --- | --- | --- |
| **Total** |  |  |  | 253.94 | 165.91 | 272.89 |
| **Paired-ends** |  |  |  | 152.78 | 115.56 | 190.07 |
|  | P01 | 180 | 100 | 74.81 | 58.35 | 95.97 |
|  | P02 | 300 | 100 | 53.82 | 39.41 | 64.82 |
|  | P03 | 500 | 100 | 24.15 | 17.80 | 29.28 |
| **Mate pair** |  |  |  | 101.16 | 50.35 | 82.82 |
|  | M01 | 2000 | 100 | 8.54 | 7.92 | 13.03 |
|  | M02 | 2000 | 125 | 12.64 | 11.67 | 19.19 |
|  | M03 | 5000 | 100 | 15.03 | 11.92 | 19.61 |
|  | M04 | 5000 | 125 | 3.82 | 3.38 | 5.56 |
|  | M05 | 10000 | 90 | 31.56 | 8.06 | 13.26 |
|  | M06 | 20000 | 90 | 29.57 | 7.40 | 12.17 |

\* The estimated genome size was ~608 Mb.

**Supplementary Table 2.** Summary of RNA sequencing (RNA-Seq) of *R. distichophylla*.

| **RNA**<br>**Source Tissues** | **Read**<br>**Length**<br>**(bp)** | **Number of**<br>**Pair-end Reads** | **Clean Data**<br>**(bp)** |
| --- | --- | --- | --- |
| **Stem** | 100 | 26,533,101 | 5,306,620,200 |
| **Root** | 100 | 27,651,149 | 5,530,229,800 |
| **Female inflorescence** | 100 | 28,030,510 | 5,606,102,000 |
| **Leaf** | 100 | 27,717,564 | 5,543,512,800 |
| **Total** |  | 109,932,324 | 21,986,464,800 |

**Supplementary Table 3.** Estimation of genome size of *R. distichophylla* based on *K*-mer analysis.

| *K*-mer | *K*-mer number | *K*-mer depth | Genome size<br>(Mb) |
| --- | --- | --- | --- |
| 17 | 57,760,011,267 | 95 | 608 |

**Supplementary Table 4.**
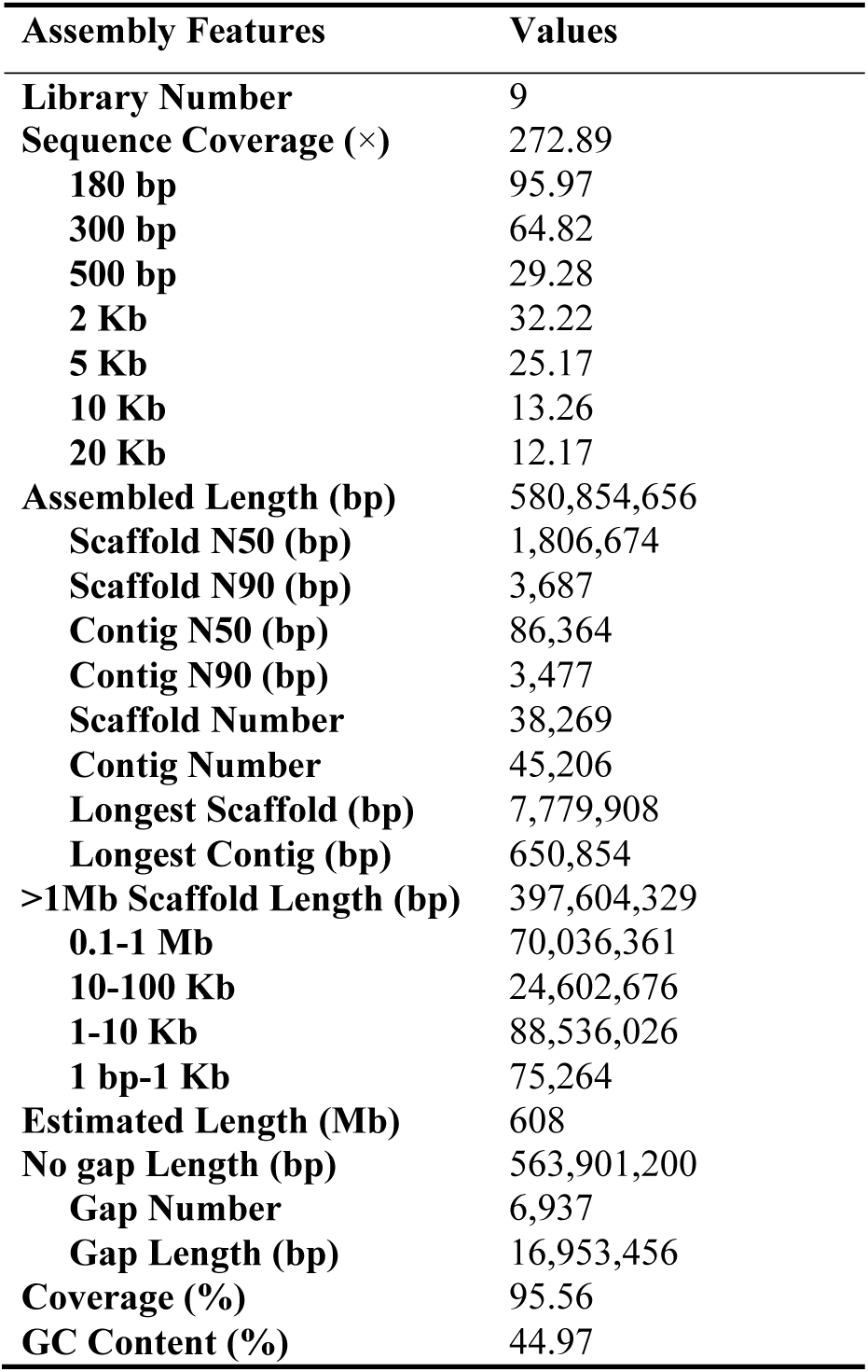
Summary of genome assembly for *R. distichophylla*.

**Supplementary Table 5.** Validation of the *R. distichophylla* genome assembly using reads mapping BUSCO, and transcript alignments.

|  | **Total** | **Mapped** | **Rate (%)** |
| --- | --- | --- | --- |
| **Reads mapping** |  |  |  |
| Short libraries | 211,270,340 | 188,569,033 | 89.25 |
| **BUSCOs from Embryophyta lineage\*** |  |  |  |
| Complete | 1,440 | 1,326 | 92.08 |
| Duplicated | 1,440 | 44 | 3.06 |
| Fragmented | 1,440 | 17 | 1.18 |
| Missing | 1,440 | 53 | 3.68 |
| **Transcripts assembled in this study\*\*** |  |  |  |
| Unigenes | 131,575 | 102,676 | 78.04 |

\* The 1,440 BUSCO conserved genes used were collected from Embryophyta lineage;
\*\* Hits with Coverage $\geq 30\%$ and Identity $\geq 80\%$ .

**Supplementary Table 6.** Statistics of repeat sequences in the *R. distichophylla* and moso bamboo genomes.

| Type | *R. distichophylla* |  | Moso bamboo |  |
| --- | --- | --- | --- | --- |
|  | Length<br>(Mb) | Percentage<br>(%) | Length<br>(Mb) | Percentage<br>(%) |
| **Total DNA TEs** | **25.78** | **4.44** | **157.02** | **7.65** |
| CMC-EnSpm | 9.19 | 1.58 | 1.54 | 0.08 |
| hAT | 5.77 | 0.99 | 50.92 | 2.48 |
| MULE-MuDR | 6.86 | 1.18 | 28.14 | 1.37 |
| PIF-Harbinger | 1.66 | 0.29 | 59.68 | 2.91 |
| TcMar-Stowaway | 1.28 | 0.22 | 6.93 | 0.34 |
| Other | 0.64 | 0.11 | 1.86 | 0.09 |
| Helitron | 0.37 | 0.06 | 7.94 | 0.39 |
| **Total No-LTR RT** | **8.73** | **1.50** | **21.97** | **1.07** |
| LINE | 7.10 | 1.22 | 21.59 | 1.05 |
| SINE | 1.63 | 0.28 | 0.38 | 0.02 |
| **Total LTR RTs** | **209.02** | **35.99** | **1014.83** | **49.46** |
| *Copia* | 47.45 | 8.17 | 344.60 | 16.80 |
| *Gypsy* | 76.77 | 13.22 | 483.77 | 23.58 |
| Other | 84.80 | 14.60 | 186.46 | 9.09 |
| **Other Repeats** | **41.57** | **7.16** | **101.85** | **4.96** |
| Satellite | 0.35 | 0.06 | 1.29 | 0.06 |
| Simple repeat | 6.04 | 1.04 | 10.50 | 0.51 |
| Low complexity | 1.12 | 0.19 | 2.03 | 0.10 |
| Unknown | 34.06 | 5.86 | 88.03 | 4.29 |
| **Total Repeat Elements** | **285.10** | **49.08** | **1295.67** | **63.15** |

**Supplementary Table 7.** Summary of types and number of simple sequence repeats in the *R. distichophylla* and moso bamboo genomes.

| **Species** | **Repeat type** | **Number** | **Proportion (%)** | **Total length (Kb)** | **Average length (bp)** |
| --- | --- | --- | --- | --- | --- |
| *R. distichophylla* | **Monomer** | 15,360 | 6.96 | 213.98 | 14 |
|  | **Dimer** | 35,195 | 15.94 | 706.01 | 20 |
|  | **Trimer** | 72,626 | 32.90 | 1063.37 | 15 |
|  | **Tetramer** | 58,046 | 26.30 | 791.80 | 14 |
|  | **Pentamer** | 26,823 | 12.15 | 442.42 | 16 |
|  | **Hexamer** | 12,687 | 5.75 | 250.85 | 20 |
|  | **Total** | **220,737** | **100** | **3468.42** | **16** |
| **Moso bamboo** | **Monomer** | 33,840 | 6.81 | 565.86 | 17 |
|  | **Dimer** | 112,350 | 22.61 | 2458.49 | 22 |
|  | **Trimer** | 129,860 | 26.14 | 1715.58 | 13 |
|  | **Tetramer** | 150,447 | 30.28 | 1963.37 | 13 |
|  | **Pentamer** | 45,721 | 9.20 | 726.52 | 16 |
|  | **Hexamer** | 24,601 | 4.95 | 505.93 | 21 |
|  | **Total** | **496,819** | **100** | **7935.75** | **16** |

**Supplementary Table 8.** Non-coding RNA genes in the *R. distichophylla* genome.

| **ncRNA Type** | **Loci#** | **Average length**<br>**(bp)** | **Total length**<br>**(bp)** | **% of genome** |
| --- | --- | --- | --- | --- |
| **miRNA** | 256 | 134.8 | 34496 | 0.0057 |
| **tRNA** | 727 | 74.3 | 53980 | 0.0089 |
| **rRNA (8S)** | 62 | 113.6 | 7043 | 0.0012 |
| **rRNA (18S)** | 12 | 1795.2 | 21542 | 0.0035 |
| **rRNA (28S)** | 16 | 4660.9 | 74574 | 0.0123 |
| **SnoRNA** | 242 | 108.3 | 26207 | 0.0043 |
| **snRNA** | 127 | 136.4 | 17314 | 0.0028 |

**Supplementary Table 9.**
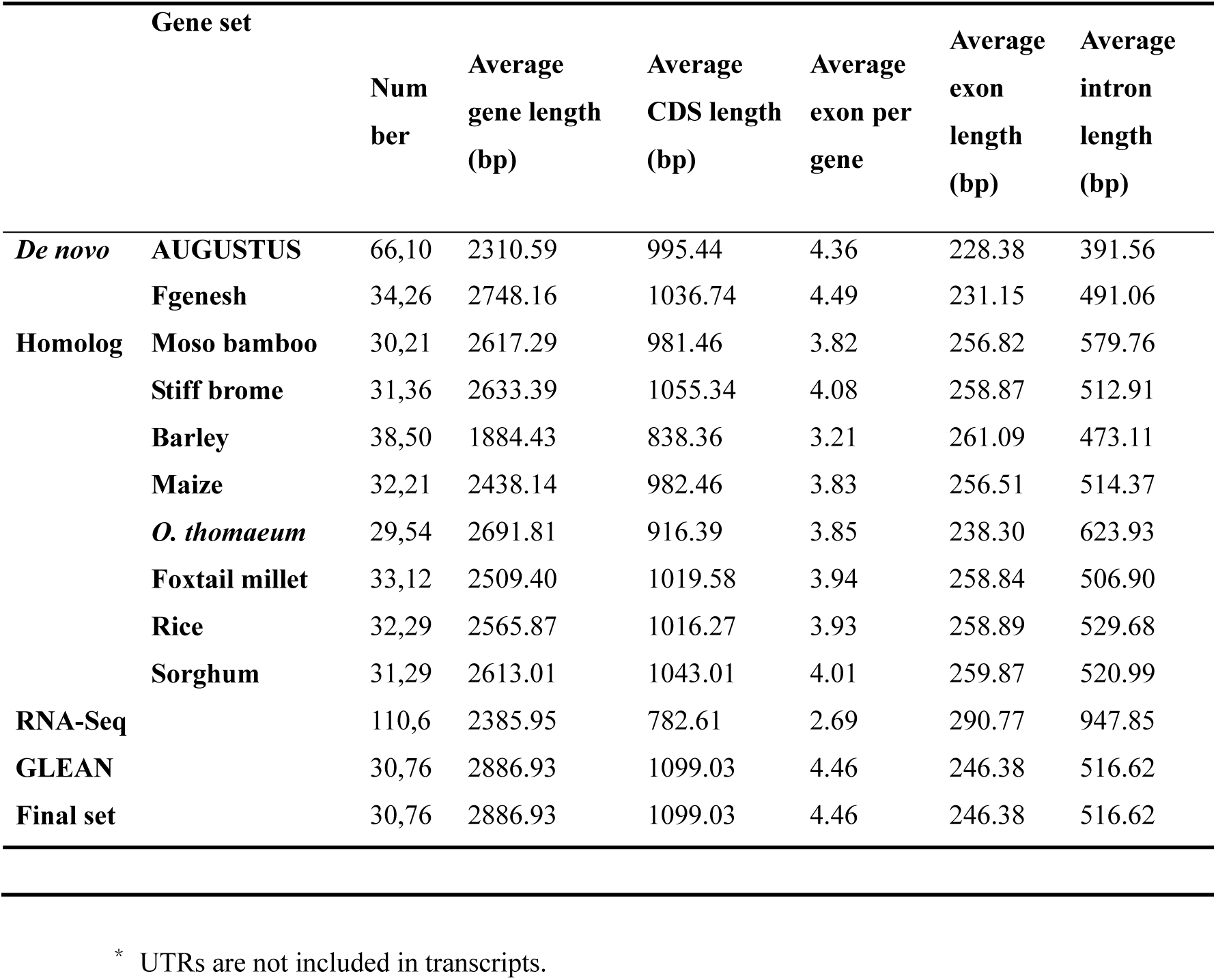
Statistics of predicted protein-coding genes in the *R. distichophylla* genome^*^.

**Supplementary Table 10.** Functional annotation of the *R. distichophylla* protein-coding genes.

| Type |  | Number | Percentage (%) |
| --- | --- | --- | --- |
| **Total** |  | 30,763 | 100 |
| **Annotated** | **InterPro** | 25,506 | 82.91 |
|  | **GO**<sup>a</sup> | 15,226 | 49.49 |
|  | **KEGG**<sup>a</sup> | 18,580 | 60.39 |
|  | **Swiss-Prot**<sup>b</sup> | 20,272 | 65.89 |
|  | **TrEMBL** | 27,191 | 88.38 |
| **Un-annotated** |  | 3,429 | 11.15 |

<sup>a</sup> Directly retrieved from InterPro entry;
<sup>b</sup> Blast hits with e-value $\leq 1e-5$ were retained.

